# Genome-Environment Associations reveal shared and unique adaptive loci across multiple pollutants and populations of the eastern mosquitofish *Gambusia holbrooki*

**DOI:** 10.64898/2026.01.07.697681

**Authors:** Antoine Fraimout, Céline Reisser, Émilie Farcy, Eva Blondeau-Bidet, Nicolas Martin, Agnes Rastteter, Arnaud Estoup, Charles Perrier

**Affiliations:** UMR CBGP, INRAE, CIRAD, IRD, Institut Agro, Université Montpellier, Montpellier, France; MARBEC, Univ Montpellier, CNRS, IFREMER, Montpellier, IRD, France; MGX-Montpellier GenomiX, BCM-BioCampus Montpellier, Université de Montpellier, CNRS, INSERM, Montpellier, France

**Keywords:** Adaptation, multi-stress, biological invasion, population genomics, eastern mosquitofish

## Abstract

Rapid adaptation to human-induced stressors is commonplace in the context of global change, including during biological invasions. Identifying the genomic bases and associated biological functions underlying such adaptation is hence crucial to understand and anticipate the response of populations and species to changing conditions. In particular, the extent to which genetic responses to multiple anthropogenic stressors vary between populations in the wild has been relatively unexplored. We addressed this question by leveraging whole-genome sequence data (both PoolSeq and IndSeq) in invasive populations of *Gambusia holbrooki* - a widespread invasive fish species - collected from 14 locations with different multi-pollutants exposure. We sought to identify adaptive loci associated with pollution tolerance by conducting Genome-Environment Association (GEA) analyses. Additionally, we investigated the degree of adaptive loci reuse between pollutants and their combinations as well as across populations. We found strong signals of association between allele frequency changes and pollutant exposure at several genomic locations, often overlapping with genes known for their functions in detoxification and immune response. We further showed that most adaptive loci are not shared among all populations, suggesting heterogeneous genomic response to each local selection. Interestingly, multiple tests looking for footprints of selection and association to pollutants yielded consistent results. Our results shed new light on the extent of genetic convergence in the genomic bases of rapid adaptation to different cocktails of human-induced pollution, and are important to our understanding of the fate of wild populations facing increasingly complex and stressful environments.

## INTRODUCTION

Global environmental changes are escalating in pace and frequency due to the intensification of human activities. As a result, natural populations face an ever-increasing number of selection pressures potentially acting on a wide range of phenotypic traits (Urban *et al*. 2023). To avoid extinction, populations must respond evolutionarily at an equally rapid pace. Our understanding of the genetic mechanisms underlying this response is central to anticipating populations’ adaptation to human-altered environments (Waldvogel *et al*. 2020). While cases of rapid adaptation to human-induced environmental change have been documented (e.g. Smith & Bernatchez 2008, Hufbauer *et al*. 2012, Papadopulos *et al*. 2021), they often involve an evolutionary response to one major environmental change exerting a strong selection pressure. For example, the rapid adaptation of insects’ populations to pesticide exposure resulting from agricultural land-use serves as a classic example of a swift evolutionary response to novel selective pressures (Hawkins *et al*. 2019). In such cases, selection is expected to favour few genes with large phenotypic effects, and mono- or oligogenic architectures are often involved in the rapid adaptation to strong selective pressures (Hansen 2006, Whiting *et al*. 2022, Höllinger *et al*. 2023).

In most cases, however, exposure to environmental pollution is more complex, with effects varying over time and ranging from chronic exposure to acute stress on organisms. Moreover, in regions where multiple human activities coexist, a diverse array of pollutants can affect ecosystems and generate complex selective pressures on populations (reviewed in Loria *et al*. 2019). The pollution of aquatic ecosystems from multiple sources (wastewater, agriculture, industry) represents such a case where multiple chemical stressors may affect aquatic organisms, and exposure to mixtures of pollutants has been reported to induce multiple behavioural, physiological and morphological alterations in both marine and freshwater organisms (Fleeger *et al*. 2003, Häder *et al*. 2020). Despite these stressful conditions, several aquatic species have evolved tolerance to pollution, and fish species in particular have become opportune models to investigate the rapid evolutionary response of aquatic vertebrates to polluted waters (*e.g*., tomcod, Wirgin *et al*. 2011; killifish, Reid *et al*. 2016; brown trout, Paris *et al*. 2024). While the evolution of tolerance to embryotoxic persistent pollutants (*e.g.,* PCBs, PAHs) seems to involve a common set of genetic pathways such as the aryl hydrocarbon receptor (AHR) pathway (Whitehead *et al*. 2012, Reed *et al*. 2016, reviewed in Whitehead *et al*. 2017), the genetic architecture underlying the tolerance to complex mixtures of sub-lethal concentrations of pollutants remains poorly understood. Overall, the evolutionary response to pollution is fundamentally multivariate, with several pollutants acting independently or in interaction on multiple, potentially correlated suites of traits.

Dissecting the genetic architecture of tolerance therefore becomes a challenge, hindered by the fact that the specific selective pressures exerted by pollutants – and the traits they target – remain elusive in the wild. In this context, population genomics approaches can complement phenotypic measurements performed in the field, by investigating the patterns of among-populations genomic variation along an environmental gradient. Specifically, changes in allele frequencies among demes occupying heterogeneous environments can be correlated to local adaptation and produce extreme values in the genome-wide *F*_ST_ distribution. Scanning the genome for such regions of high differentiation can inform on the loci under selection in a specific environment (Lotterhos & Whitlock 2015). Nonetheless, disentangling whether genomic regions exhibit differentiation between populations because of local adaptation, or chance events is in turn a difficult task, as neutral evolutionary processes may well generate patterns of high *F*_ST_ values and false-positive signals of local adaptation (Gautier 2015). In the case of invasive species in particular, complex demographic histories involving founder effects, genetic bottlenecks or geographic isolation can generate strong signals of differentiation in the genome due to an increase of genetic drift in small populations. Understanding the genomic bases of adaptation to complex patterns of pollution and the evolution of tolerance thus requires considering both the diverse environmental pressures exerted by multiple pollutants, and the complex evolutionary histories of the affected populations.

This context raises several broad questions that could improve our understanding of adaptation to human-modified environments and rapid evolution in natural populations. A first question concerns the pace at which genetically based adaptation can emerge under exposure to multiple pollutants. Most contaminants occur at sub-lethal levels and typically induce only mild phenotypic effects, making strong selection and rapid allele-frequency shifts unlikely (Aich *et al*. 2024). Yet, even moderate pollution can reduce fitness - through declines in fecundity, compromised physiological performance, or indirect ecological effects such as altered resource availability (Saaristo *et al*. 2018). A second question concerns the dimensionality of genetic adaptation to pollutant cocktails. Multiple contaminants impose multiple selection pressures, often acting on several traits and it remains unclear whether the genetic architecture underlying tolerance scales with the complexity of these selection factors. A third related question is whether pollution tolerance, despite being shaped by diverse contaminants, relies on conserved genetic bases or instead on genetic redundancy, where different loci can generate similar phenotypic responses (Láruson *et al*. 2020).

Here, we used populations of the eastern mosquitofish, *Gambusia holbrooki,* that recently invaded Europe (Pyke 2005, 2008) as a study model to address these questions. *G. holbrooki* is a small live-bearing teleost fish species from the *Poeciliidae* family, native from continental USA (Pyke 2005). In the last century, it has been used as a biological control agent against mosquito larvae and deliberately introduced in dozens of countries across all continents except Antarctica (Pyke 2008). As one of the most invasive fish species worldwide, *G. holbrooki* is characterized by its large population sizes and relatively short generation time (ca. four weeks; Fernandez-Delgado 1989, Vila-Gispert & Moreno-Amich 2002), and its tolerance to a large range of environmental conditions (Pyke 2005). In Europe, *G. holbrooki* was first introduced in 1921 (Krumholz 1948) and is now present in most Mediterranean countries, where it is commonly found in freshwater habitats such as ponds and rivers, but also in brackish, and even saltwater environment in littoral areas (Alcaraz & Garcia-Berthou 2007, current study). Invasive populations of *G. holbrooki* are also well known for their capacity to tolerate highly polluted environments. For example, Díez-del-Molino *et al*. (2018) studied invasive *G. holbrooki* populations in the Ebro River in Spain, an ecosystem experiencing high levels of chemical disturbance from various pollutants (*i.e.,* heavy metals and organochlorine pesticides), and found some evidence of genetic differentiation associated with tolerance to pollution based on microsatellite and candidate gene markers. The recent European introduction of *G. holbrooki* thus provides an ideal system for investigating the genetic basis of rapid adaptation to multiple chemical stressors in natural environments.

Using environmental data (*i.e*. pollution profiles) and whole-genome sequencing data from 14 Mediterranean populations of *G. holbrooki*, we examined how genomic variation tracks exposure to complex mixtures of contaminants. Together, these data allow us to assess the pace and mechanisms of genetic adaptation to human-altered environments shaped by multiple, simultaneous stressors. We more specifically addressed two questions: (i) do populations exposed to different pollution sources show genomic signatures of local adaptation? and (ii) what is the genetic architecture underlying tolerance to mixtures of contaminants? More specifically, we focused on identifying the genes and biological pathways associated with these contaminant regimes, and on assessing whether genomic responses are shared or distinct across populations experiencing different pollutant regimes.

## MATERIAL AND METHODS

### Population sampling, DNA extraction and resequencing

Sampling of *G. holbrooki* populations is described in detail in Martin *et al*. (2025a). To summarize, 14 locations were sampled across a ca. 150 km area (Fig. 1A, Table S1) in Southern France, corresponding to documented gradients of industrial, agricultural and urban wastewater pollution overlapping with mosquitofish distribution in the area. Genomic DNA was extracted using a Maxwell® 16-LEV Blood DNA Kit. We then prepared DNA for: i) individual sequencing of 2-10 fish per population (hereafter “IndSeq” dataset; see Table S1) and ii) one pool sequencing of 30 individuals (15 males and 15 females) per population (hereafter “PoolSeq” dataset). All samples were sequenced at the Montpellier Genomix (MGX) platform.

**Figure 1.**
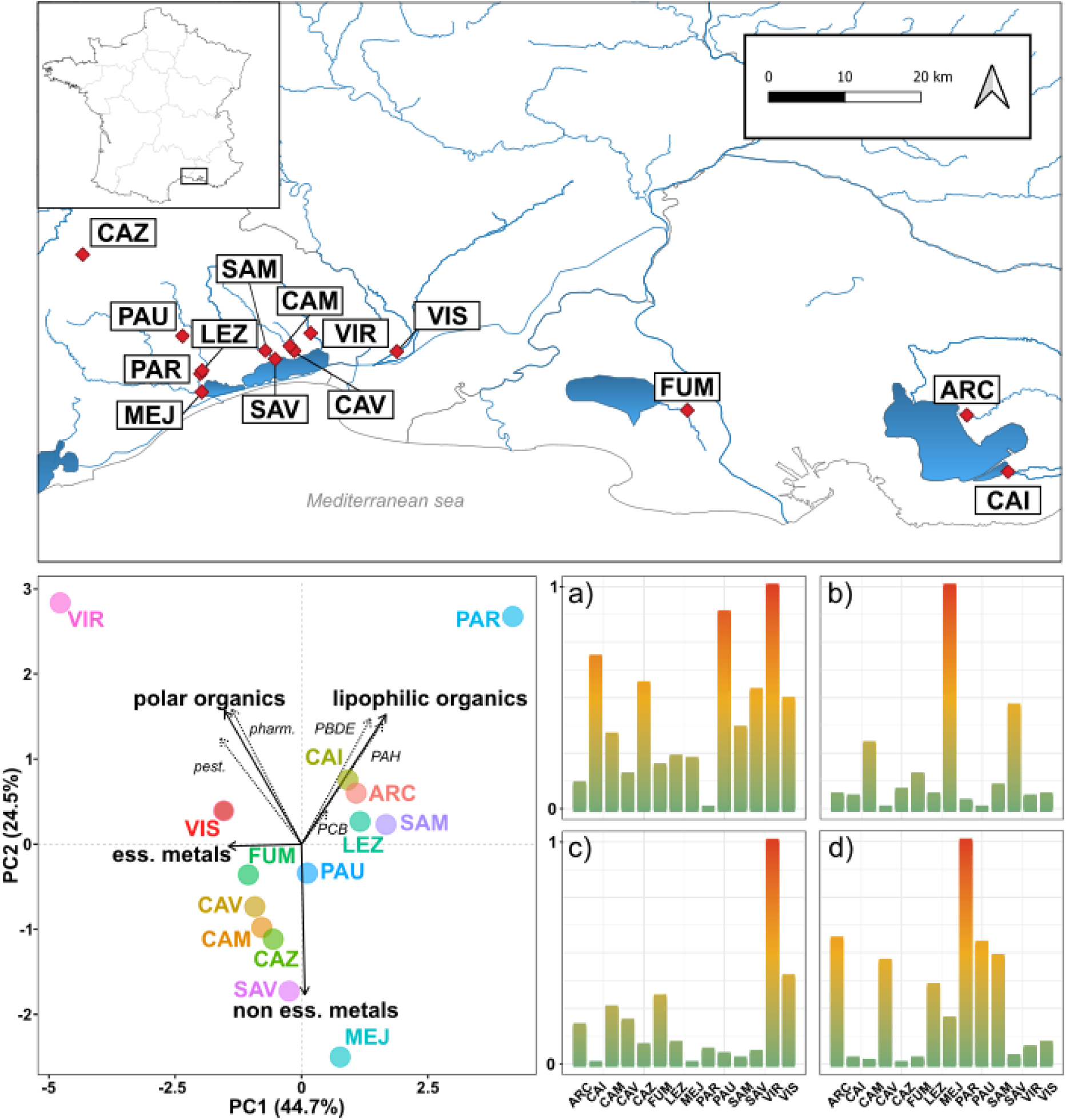
Sampling and characterization of environmental pollution. The sampling locations of our study populations are shown on the map (top panel). The results of the PCA on pollution measurements (bottom-left panel; see *Methods* for details) show the four main pollution variables (bold arrows and text) used for the GEA. Histograms of scaled pollution measurements (four bottom-right panels) are shown for all populations and colored along a gradient from low (green) to high (red) values for a) essential metals, b) non-essential metals, c) polar organics and d) lipophilic organics.

### Characterization of environmental pollution

We used the datasets from Martin *et al*. (2025a, b) to characterize the pollution landscape of our study populations. Briefly (but see detailed methods in Martin *et al*. 2025a, b), pollutants were measured in three different media: water column, whole fish tissue and muscle tissue. From the water column, Polar Organic Chemical Integrative Samplers (POCIS) were used to measure the concentration of pesticides and pharmaceutical compounds. Persistent organic pollutants (*i.e.,* polycyclic aromatic hydrocarbons [PAHs], polychlorinated biphenyls [PCBs] and polybrominated diphenyl ethers [PBDEs]) were measured from whole fish tissue using a combination of solvent extraction and gas chromatography, and were normalized from the total lipid content. Finally, the concentration of essential (Zn, Cu, Ni, Cr) and non-essential (As, Cd, Pb, Hg) metals was assessed from muscle tissues using mass spectrophotometry.

Prior to conducting the GEA, we performed a principal component analysis (PCA) of the raw pollution dataset (Fig. 1), using all scaled and centered variables and the *prcomp* function in R (v. 4.4.2; R Core Team 2024).

### SNP calling

BWA-MEM (Vasimuddin *et al*. 2019) was used to align raw reads on the reference genome assembly (Hap1) from a local invasive population from the PAU population in Montpellier (Cormier *et al*. in prep; genome data available at https://bioinfo.gitlab-pages.ifremer.fr/bioanalysis/internal/gamboc/) to generate bam files. Variant detection was performed with the Bayesian software freebayes (Garrison & Marth 2012). We filtered variants based on mapping and base quality (options *-m 20* and *-q 20*, respectively) and applied the pooled-continuous option to jointly analyse the pooled and individual sequencing datasets (hereafter PoolSeq and IndSed datasets). We also limited coverage range from 122 to 250 using the *min-coverage* and *limit-coverage* options, respectively, and used the *hwe-prior-off* option to disable estimation of Hardy-Weinberg Equilibrium probabilities. The VCF output was further filtered using bcftools (*QUAL>20 & TYPE=“SNP” & SAF > 0 & SAR > 0 & RPR > 0 & RPL > 0*). After filtering, the resulting dataset comprised 3,124,724 biallelic SNPs across 14 populations from which we created two VCF files for the PoolSeq data and the IndSeq data.

### Characterization of genetic diversity and structure between populations

We used the *poolfstats* R package (v.3.0.0; Gautier *et al*. 2022) to characterize the genomic differentiation within and among populations of the PoolSeq data. We used the *min/max.cov.per.pool* (set to 20 - 180) options of the *vcf2pooldata* function in *poolfstats* to further filter the data based on the minimum/maximum read count per pool. The resulting filtered PoolSeq data consisted of 2,877,126 biallelic SNPs. To describe patterns of genomic variation across populations, we estimated genome-wide heterozygosities within populations using the *compute.fstats* function and calculated pairwise *F*_ST_ between populations using the *compute.pairwiseFST* function. We also calculated the pairwise divergence index (*dxy*) using the *pixy* software (v.1.2.11; Korunes & Samuk 2021).

### Genome-Environment Association analysis

We performed whole-genome scans for signals of putative local response to selection and of putative response to selection induced by pollutant exposure using BayPass v.2.4 (Gautier 2015). First, we ran the core model of BayPass to estimate Ω, the scaled covariance matrix of population allele frequencies. The PoolSeq dataset for all 14 populations was sub-sampled into 38 data files each containing 75,000 SNPs, using the *pooldata2genobaypass* R function. We ran the BayPass core model on each data subset using default Markov Chain Monte Carlo parameters (MCMC; 5000 burning, thinning every 20th iteration). We assessed model convergence by computing the pairwise distance between Ω generated from three independent chains by adjusting the *seed* option in BayPass. Specifically, we used the *fmd.dist* R function implemented in BayPass to compute the pairwise FMD distances (Förstner & Moonen 2003) between all Ω matrices within each run (among all 38 matrices per run) and among runs. Given the narrow distributions of both between- (mean FMD 0.006 ± 0.001) and within-run distances (mean FMD 0.15 ± 0.002), we retained a single Ω matrix (computed from the first subsample or the first run) for further analyses.

We then tested for associations between SNPs and pollutant exposure using the auxiliary model in BayPass. We parametrized the model using Ω with the *–omega* argument to account for the effect of potentially complex demographic histories in our data, and ran it with default MCMC parameters, using the –*auxmodel* argument. Based on the variation in pollution exposure among populations from the PCA plot (Fig. 1), we used four main variables to describe pollutant exposure: i) the concentration of lipophilic organics, measured in the whole tissues of fish and representative of the body burden of exposure to POPs; ii) the concentration of polar organics (pharmaceutical and pesticide compounds) measured in the water column at each samples site; iii) and iv) the muscle tissue concentration of essential and non-essential metals, respectively. We thus fitted four separated auxiliary models including each main pollution variable as an environmental covariate and ran each model in parallel across the 38 data subsets.

BayPass outputs were aggregated using the *concatenate_res* R function and analysed in R. Computation of Ω with the core model allowed obtaining the XtX statistic of genomic differentiation at all SNP positions (Günther & Coop 2013). The XtX is defined as the variance of the standardized population allele frequencies and provides a measure of SNP differentiation, while accounting for covariance structure in population allele frequencies (Günther & Coop 2013, Gautier 2015). In other words, XtX measures highly differentiated SNP among populations while correcting for the confounding effects of genetic drift or demographic history. From the auxiliary model outputs, significant association between a SNP and a pollution variable was determined following BayPass best practices (Gautier 2015) by considering all SNPs with a Bayes Factor (BF) value above 30. Finally, we used the local-score approach (Fariello *et al*. 2017) to delineate windows of significant genomic differentiation or association with pollution exposure from the core and auxiliary model outputs, respectively. This approach allows defining genomic windows with an excess of highly significant SNP (*i.e.* with high BF values or low *p*-values for the auxiliary and core model, respectively) showing association with each pollution variable. To this end, we used the *compute.local.scores* R function available in BayPass and set the ξ parameter to 2.

### Comparison of nucleotide diversity between genomic regions

We estimated genome-wide nucleotide diversity (π) within each population using *pixy* (Korunes & Samuk 2021). More specifically, we estimated and compared the levels of nucleotide diversity across four types of genomic regions in our data: (i) “XtX”, corresponding to the genomic regions where highly differentiated windows were delineated by the local-score approach from the *p-values* of the XtX statistic; (ii) “pollution”, corresponding to all the windows significantly associated with each type of pollutant (based on the local-score) and (iii) ‘background’, corresponding to regions showing neither XtX signals nor pollution-associated signals of selection. We also compared the overall nucleotide diversity in adaptive genomic regions (*i.e.* across all XtX significant windows) *vs.* background genomic regions. We followed standard practice for the *pixy* software (https://pixy.readthedocs.io/) and estimated π in 10kB windows across the genome. Results from *pixy* were analysed in R: we used the *GenomicRanges* package (v.1.58.0, Lawrence *et al*. 2013) to identify overlaps between the adaptive windows and each 10kB window delineated by *pixy* to calculate the average π in adaptive genomic regions. Average π for background genomic regions was calculated on those regions not overlapping with any adaptive genomic regions.

### Haplotype-based search for genomic footprints of selection

We used a different and complementary method to scan the genome for signals of selection by running on the IndSeq dataset the haplotype-based approach implemented in the *rehh* R package (v.3.2.2; Gautier *et al*. 2017). Under positive selection, a beneficial allele will rise in frequency in the population and recombination in the region around that allele will be reduced, thus generating stretches of homozygosity. By measuring the decay of haplotype homozygosity around a favourable allele, Sabeti *et al*. (2002) proposed the concept of Extended Haplotype Homozygosity (EHH) as a test to detect footprint of positive selection. Here, we used the *rehh* package to estimate the between-population XP-EHH summary statistic, and identify differential signal of positive selection between pairs of polluted and non-polluted populations along our four main categories of pollutants. Prior to running this analysis, we used Beagle (v.4.0; Browning & Browning 2013) on the IndSeq dataset to phase the data for each population separately. We then imported the phased data in *rehh* and used the *ies2xpehh* function for pairs of populations of interest. As our measure of pollution exposure is continuous and populations are not classified in discrete pollution groups, we defined pairs of populations by partitioning the data into clusters of polluted *vs.* non-polluted populations. We used the *pam* function from the *cluster* R package (v.2.1.8.1; Maechler *et al*. 2013) on each vector of continuous pollution values and set the number of clusters to three to obtain clusters of low, medium and high pollution values. Pairs of populations were then constituted by choosing the populations from the low- and high-pollution clusters and we calculated the XP-EHH statistics for all these pairs, for each type of pollutant (*i.e.,* lipophilic organics, polar organics, essential and non-essential metals, as defined in the *Genome-Environment Analysis* section). We used the *calc_candidate_regions* function to delineate genomic regions containing clusters of outlier markers, as measured by their strongly negative *p-*value associated with the XP-EHH. We replicated the *rehh* analysis on filtered data by discarding low frequency variants (*i.e.,* those with a MAF < 0.05 in at least one population) using plink (v.1.9, Purcell *et al*. 2007).

### Gene Ontology (GO) enrichment analysis

We identified the genes contained in all significant windows using the *bed_intersect* function of the *valr* R package (v.0.8.2; Riemondy *et al*. 2017). For each type of pollutant, we produced a list of genes from the overlap between the gene annotation file (GFF) and the pollution windows delineated with the local-score approach. We then performed a test of GO term enrichment to highlight the biological processes enriched amongst the genes with pollution-associated SNPs. We gathered GO terms for our annotated genome from UniProt (https://www.uniprot.org/) through a batch search of all protein names from our annotated GFF. This resulted in 7,735 matches with 3,141 GO terms for *Biological process*, which we used as the background against which to contrast our pollution-associated genes. We used the *topGO* R package (Alexa & Rahnenfuhrer 2009) to perform exact tests of enrichments and restricted our analysis to genes within a 5kb distance from the pollution-associated genomic windows.

## RESULTS

### Pollution exposure and genomic variation among populations

We observed wide variation in pollutant exposure among the sampled sites (Fig. 1). While some populations showed low concentration levels for all types of pollutants, other populations were exposed to high levels of a specific type of pollutant (*e.g.* PAR to polar organics; Fig. 1) or to medium levels of multiple types of pollutants (*e.g.*, SAM to metals). Genetic differentiation among populations was rather modest: the first two PCA axes explained 17.71% of the variance (Fig. 2A), and the median genome-wide *F*_ST_ was 0.099 (Fig. 2B, Fig. S1). Two populations (PAU and SAM) stood out as more differentiated from all others and from each other. Aside from these, population genetic structure was shallow. Genome-wide *dxy* values ranged from 2.024 × 10⁻³ (CAV–SAV) to 2.346 × 10⁻³ (CAZ–SAM; Fig. 2B) and highlighted geographic patterns of divergence, with the isolated CAZ population showing greater divergence from the rest.

**Figure 2.**
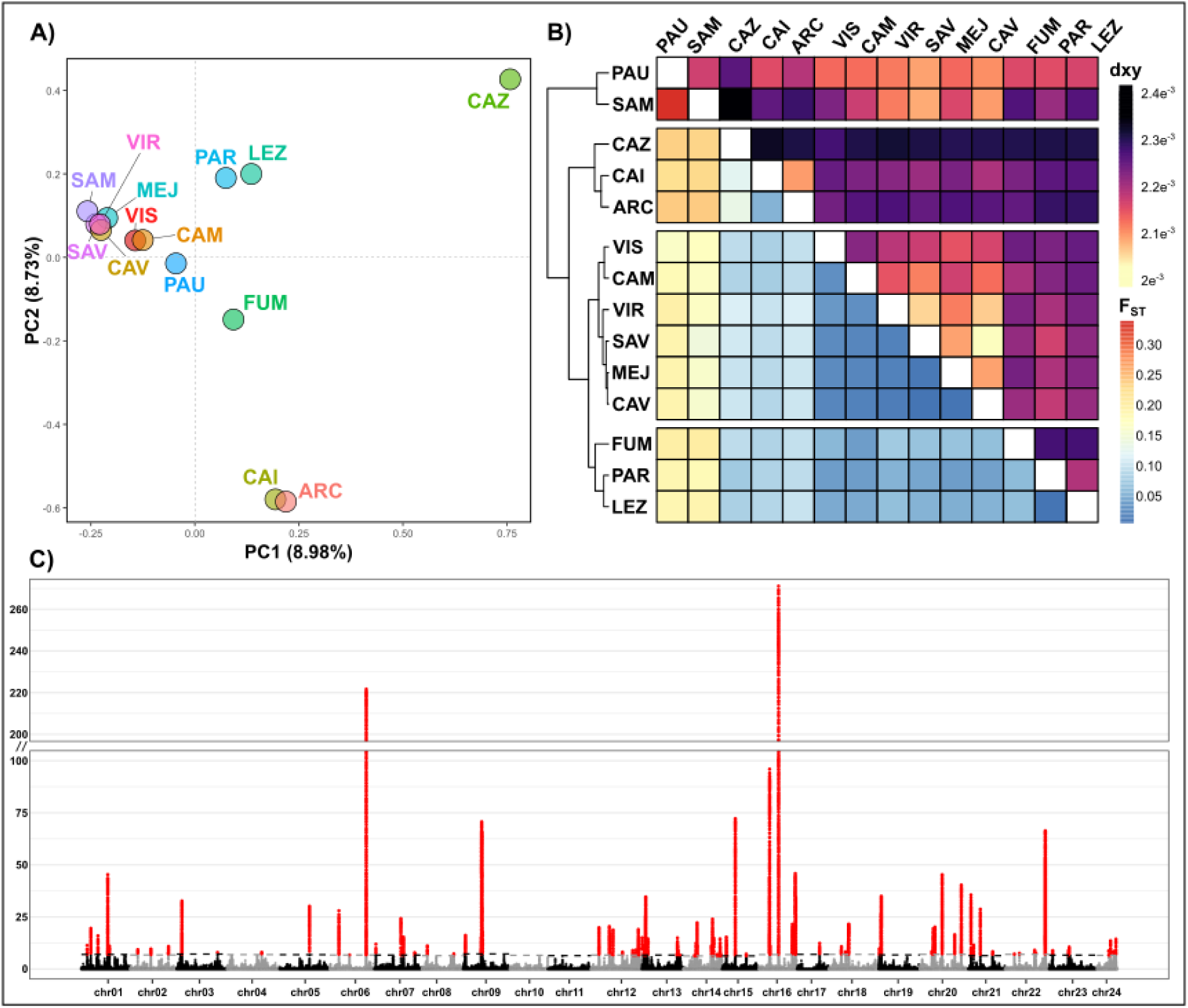
Genome-wide variation and signal of adaptation. A) Principal Component Analysis of genomic variation among populations. B) The pairwise F_ST_ matrix (lower triangle) between all pairs of populations is shown. Colour gradient represents low (blue cells) to high (red cells) F_ST_ values. The dendrogram was constructed from the pairwise F_ST_ matrix to represent genetic distances between populations. The pairwise dxy matrix (upper triangle) between all pairs of populations is shown. Colour gradient represents low (yellow) to high (black) dxy values. C) Manhattan plot of the SNPs significantly associated (-log10(p) > 4; red dots) within the genomic windows delineated by the local-score (y-axis) approach run on the BayPass core model outputs. Parallel lines on the y-axis indicate axis break.

**Figure 2.**
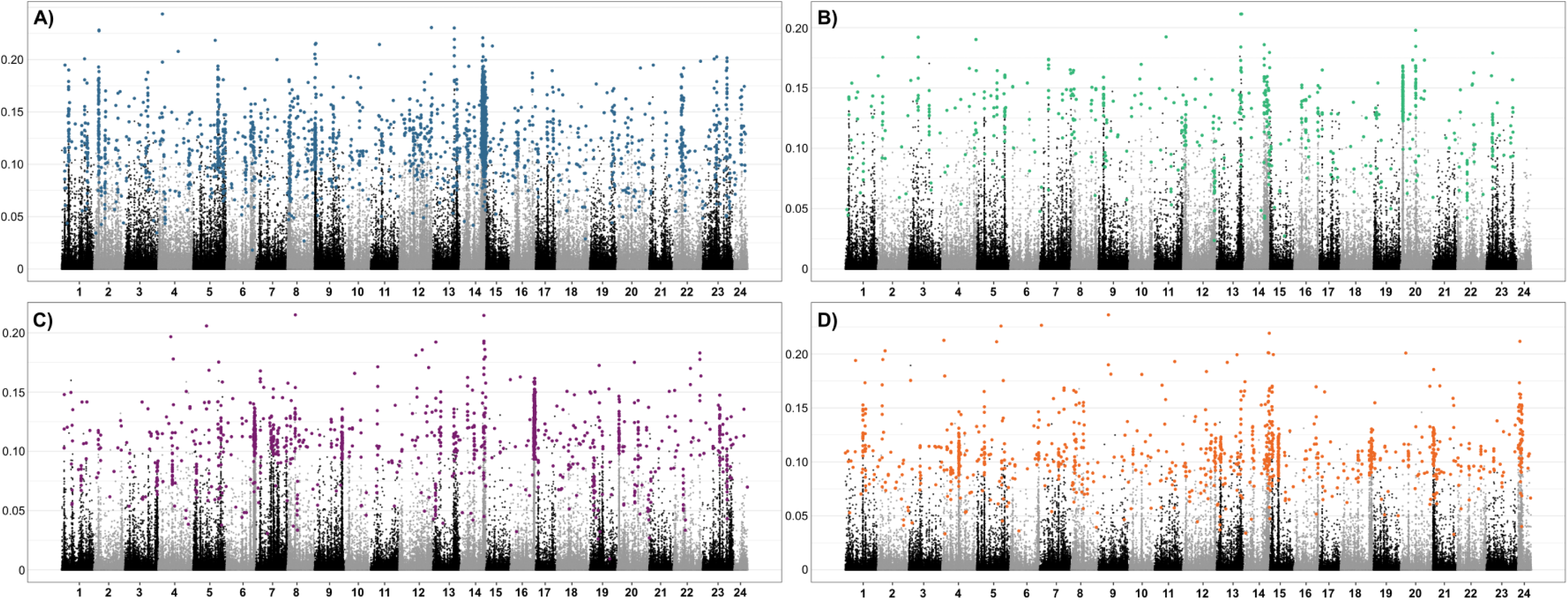
Results from the GEA analyses. For each type of pollutant (A - D) the Manhattan plot shows the SNPs significantly associated (BF > 30; colored dots) along with the coefficient of regression (β, y axis) from the BayPass auxiliary model. A: Lipophilic organics; B: Polar organics; C: Essential metals: D: Non-essential metals

### Genomic signals of positive selection and adaptation to pollutants

#### a) Genome-wide signal of adaptation

We found 5,789 SNPs across all 24 chromosomes showing a significant *p-value* (-log10(P*) >* 3) associated with the XtX statistic, and therefore putatively under positive selection (Fig. S4). The local-score approach (Fariello *et al*. 2017) delineated 162 significant windows containing a total of 9,190 SNPs distributed across almost all chromosomes (chromosome 10 and 11 only did not contain any significant windows; Fig. 2C). Windows with the highest local score values (*i.e.* the strongest ‘peak’) were located on chromosome 6 and chromosome 16 (Fig. 2C).

#### b) Association with lipophilic organics (PCB, HAP, PBDE)

Results from the BayPass auxiliary model indicated that 2,645 SNPs across all 24 chromosomes were significantly associated (BF value > 30) with exposure to lipophilic organics. Significant SNPs were also found to have strong effects as indicated by the regression coefficient obtained from the auxiliary model (Fig. 3A). The local-score approach identified 1,745 SNPs distributed in 62 genomic windows across eight chromosomes showing significant association with exposure to lipophilic pollutants (Fig. 3B). The majority of these SNPs (1,416 SNPs, representing 81.15%) were located on chromosome 14, while the remaining SNPs were found on chromosomes 2, 4, 5, 6, 9, 13 and 16 (Fig. 3A). Windows with the highest local score values were located on chromosome 14.

**Figure 3.**
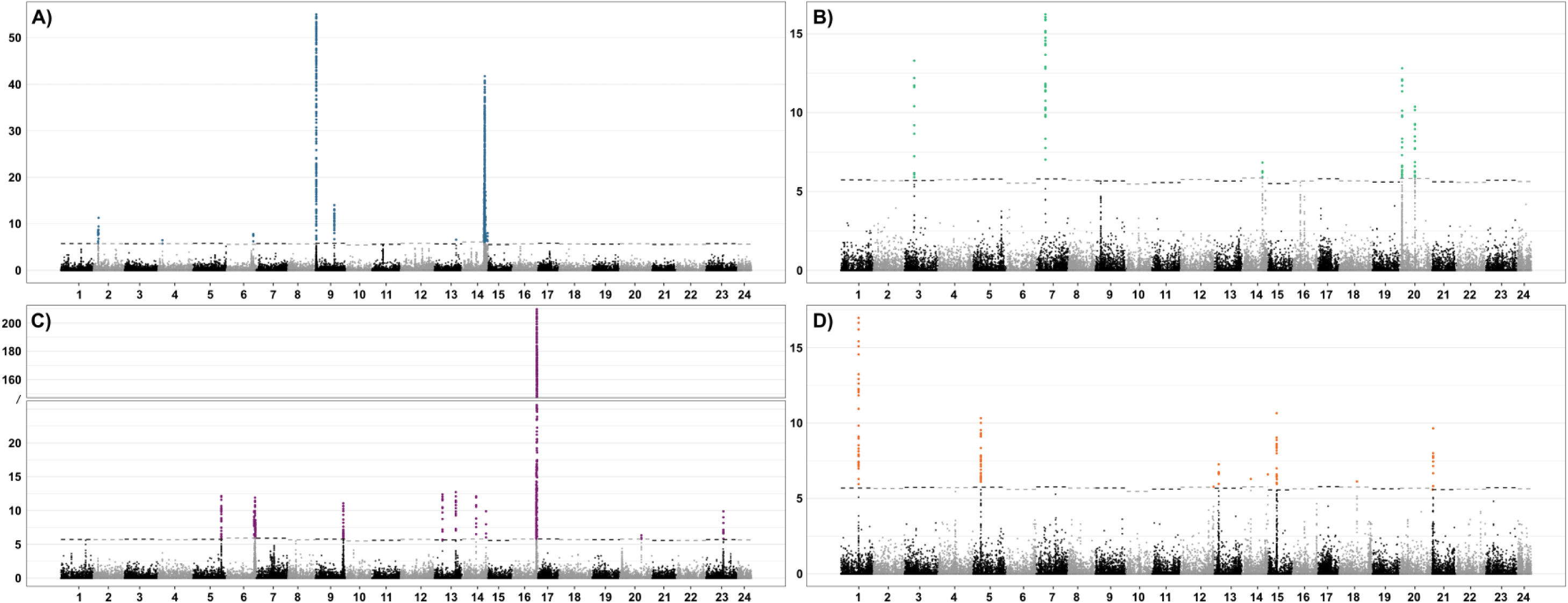
Results from the local-score approach. For each type of pollutant (A - D) the Manhattan plot shows the SNPs significantly associated (-log10(p) > 4; colored dots) within the genomic windows delineated by the local-score (y-axis) approach ran on the BayPass auxiliary model outputs. A: Lipophilic organics; B: Polar organics; C: Essential metals: D: Non-essential metals. In C) the slash symbol indicates axis break.

#### c) Association with polar organics (pharmaceutical and pesticide compounds)

We found 1,291 significant SNPs associated with exposure to polar organics (Fig. 3C) across all 24 chromosomes and 1,246 SNPs in 28 genomic windows across 10 chromosomes following the local-score approach. There was a strong and neat peak located on chromosome 16 corresponding to a genomic window containing 60.27% of all significant SNPs (751 SNPs; Fig 3D) from this association. Other significant peaks of association were located on chromosomes 1, 5, 6, 8, 9, 13, 14, 20 and 23.

#### d) Association with essential and non-essential metals

There were 16 (312 SNPs) and 19 (296 SNPs) genomic windows associated with essential and non-essential metals, respectively. Despite the similar number of windows between the two categories, significant SNPs for essential metals were located across 6 chromosomes (3, 7, 9, 14 and 20) and across 12 chromosomes for non-essential metals (Figure 3D).

### Nucleotide diversity in adaptive vs. background genomic regions

Overall, we found that genomic regions showing signals of positive selection (*i.e.* regions containing significant windows associated with the XtX statistics, or associated with a type of pollutant) showed higher nucleotide diversity than background genomic regions across all populations (Fig. S2; ANOVA, F = 5371.643, *p <* 2e-16). Furthermore, we found that both categories of adaptive regions (*i.e.* XtX- or pollution-associated) had higher diversity than background regions. Finally, we found that the genomic regions associated with lipophilic organics had higher diversity than all other adaptive regions (*p <* 2e-16 across all populations; Fig. S2).

### Haplotype-based evidence for selection

We further assessed the robustness of the signals of selection from the PoolSeq-based BayPass analysis by investigating the overlap between the significant pollution-associated windows and the XP-EHH candidate regions delineated by *rehh* analysis on IndSeq data (Table 1). There was a good concordance between the significant genomic regions identified by both approaches, across multiple pairs of populations. In other words, regions of the genome identified by BayPass to be associated with pollution exposure also showed evidence for selection based on the pairwise tests contrasting polluted and non-polluted populations. Across population pairs, overlap between local-score and XP-EHH windows averaged 78.12% for lipophilic organics, 74.01% for polar organics, 43.30% for essential metals, and 51.67% for non-essential metals (Table 1).

**Table 1.** Results of the XP-EHH test of selection. i) Pollutant: the type of pollutant used for the BayPass association analyses; ii) No. of LS windows: the number of genomic windows delineated by the local-score approach performed on the results of the BayPass association analyses. iii) Population pair: the pairs of populations corresponding, for each type of pollutant, to the most polluted (left-hand side) and the least polluted (right-hand side) population. iv) No. of LS windows included in XP-EHH candidate regions: the number and percentage of overlaps between genomic windows delineated by the local-score approach and the candidate regions of significant XP-EHH statistics.

| Pollutant | No. of LS windows | Population pair<br>(polluted vs. non-polluted) | No. of LS windows<br>included in XP-EHH candidate<br>regions |
| --- | --- | --- | --- |
| Lipophilic organics | 62 | PAR - VIS | 29 (46.77%) |
|  |  | PAR - MEJ | 61 (98.39%) |
|  |  | PAR - FUM | 59 (95.16%) |
|  |  | PAR - SAV | 11 (17.74%) |
|  |  | PAR - CAV | 55 (88.71%) |
|  |  | PAR - CAM | 54 (87.10%) |
|  |  | PAR - CAZ | 58 (93.55%) |
|  |  | PAR - VIR | 60 (96.77%) |
| Polar organics | 28 | VIR - PAU | 9 (32.14%) |
|  |  | VIR - PAR | 18 (64.29%) |
|  |  | VIR - MEJ | 24 (85.71%) |
|  |  | VIR - SAV | 19 (67.86%) |
|  |  | VIR - CAM | 25 (89.29%) |
|  |  | VIR - SAM | 27 (96.43%) |
|  |  | VIR - CAZ | 21 (75%) |
|  |  | VIR - LEZ | 21 (75%) |
| Essential Metals | 16 | PAU - PAR | 7 (43.75%) |
|  |  | PAU - MEJ | 5 (31.25%) |
|  |  | PAU - FUM | 7 (43.75%) |
|  |  | PAU - CAV | 4 (25%) |
|  |  | PAU - CAI | 6 (37.5%) |
|  |  | PAU - ARC | 1 (6.25%) |
|  |  | PAU - LEZ | 8 (50%) |
|  |  | VIR - PAR | 14 (87.5%) |
|  |  | VIR - MEJ | 16 (100%) |
|  |  | VIR - FUM | 13 (81.25%) |
|  |  | VIR - CAV | 15 (93.75%) |
|  |  | VIR - CAI | 11 (68.75%) |
|  |  | VIR - ARC | 7 (43.75%) |
|  |  | VIR - LEZ | 13 (81.25%) |
| Non-Essential Metals | 19 | MEJ - VIS | 11 (57.89%) |
|  |  | MEJ - PAU | 2 (10.53%) |
|  |  | MEJ - PAR | 8 (42.11%) |
|  |  | MEJ - FUM | 6 (31.58%) |
|  |  | MEJ - CAM | 8 (42.11%) |
|  |  | MEJ - SAM | 6 (31.58%) |
|  |  | MEJ - CAZ | 6 (31.58%) |
|  |  | MEJ - CAI | 6 (31.58%) |
|  |  | MEJ - ARC | 6 (31.58%) |
|  |  | MEJ - VIR | 10 (52.63%) |
|  |  | MEJ - LEZ | 10 (21.05%) |

### GO enrichment analysis

For each type of pollution, we identified several GO biological process terms significantly enriched in annotated genes whose sequences are overlapping with the genomic windows delineated by the BayPass/PoolSeq/local-score approach (Fisher’s test; *p* < 0.05, Table S2). A total of 14 biological processes were enriched within genes associated with exposure to lipophilic organics (Table S2). The top enriched GO term for this type of pollutant was “innate immune response” (GO:0045087). Six biological processes were also related to immunity including regulations of interferon production (GO:0032727/0032728 and 0060337), inflammatory response (GO:0050727 and GO:0050728) and antiviral immune response (GO:0140374). Genes associated with exposure to polar organics were significantly enriched in 22 biological processes, with the two top GO terms involved in cholesterol metabolic processes (GO:0008203) and negative regulation of BMP signaling pathways (GO:0030514). For metal exposure, we found two processes related to embryogenesis significantly enriched in genes associated with essential metals (GO:0048598 and GO:0009790), and 12 processes enriched in genes associated with non-essential metals, with the majority related to fatty acids or lipid metabolism (Table S2).

## DISCUSSION

We analysed *G. holbrooki* populations originating from a heterogeneous landscape of pollution exposure, where a few populations are exposed to relatively high concentrations of specific pollutants (*e.g.*, VIR to polar organics, PAR to lipophilic organics), and other populations are either exposed to medium levels of multiple pollution or consistently found in a low pollution area (Fig. 1). Our results show that rapid adaptation to such multiple human-induced stress can unfold in invasive populations despite reduced standing genetic variation. More specifically, our GEA analyses allowed identifying genomic regions putatively involved in adaptation to pollutants exposure. These adaptive loci appeared to be specific to each type of pollution, suggesting distinct polygenic genomic architectures evolved in response to different pollution exposure. In the following, we discuss our results, their implications and the potential shortcomings of our study.

### A mild and heterogeneous selective landscape of pollution

Although the analysed *G. holbrooki* populations were exposed to multiple pollutants, we found that the concentrations of the different types of pollutant measured in the water, or directly in the fish, were relatively mild in comparison to heavily polluted sites in other systems (*e.g.* concentrations of PAHs measured in Reid *et al*. 2016). This suggests that the selective effect of pollution exposure - *i.e.* the lethality or reprotoxicity of the pollutants - is unlikely to be strong, especially for a pollutant-tolerant species like *G. holbrooki*. Multiple sources of pollutants at the concentrations measured in our study are indeed unlikely to pose an acute toxicity and induce direct mortality (Whitehead 2017). Instead, they are expected to affect various physiological processes without imposing mortality-level stress. The selective landscape of pollution is thus likely heterogeneous, mostly including indirect effects on behaviour, immunity or fecundity (Scott & Sloman 2004, Segner *et al*. 2012, Hamilton *et al*. 2017, Jacquin *et al*. 2020) and indirect effects through other environmental parameters such as resource availability and quality (*e.g.,* accumulation of metal pollutants in lower trophic levels; Barrett *et al*. 2018). Another important aspect of selection by pollution is the temporal dynamics of pollutants exposure, which we could not address with the data at hand. Specifically, estimating whether populations are subjected to chronic and steady levels of pollution, or bouts of strong exposure would be valuable to assess the mode of selection (*e.g.,* balancing selection) imposed by pollution.

### Genetic bases of adaptation to complex contaminant regimes

Dissecting the genomic regions located under the peaks of association with pollutants, we found several well-characterized genes and biological processes underlying the response to each type of pollutant. We found a prevalence of immunity-related genes associated with exposure to lipophilic organics and found that immunity was the most significant biological process enriched for these genes. Particularly, the tripartite motif (TRIM) protein-coding families are known for their role in the interferon pathway in several animal species, including fish (Langevin *et al*. 2019), and are involved in mediating pathogens recognition and immune responses to microbial and viral infections (Ozato *et al*. 2008). This suggests that the adaptive signal we detect in our study likely involves selection acting on variants conferring advantageous immunity to individuals exposed to such pollution. We cannot rule out that populations exposed to lipophilic organics are not also exposed to a high pathogen load, which could in turn explain the selection for immune response. However, the effects of lipophilic organics on fish immune response have been described in several fish species (Reynaud & Deschaux 2006). Particularly, exposure to polycyclic aromatic hydrocarbons (PAH; a constituent of our lipophilic organics category of pollutant) has been shown to decrease host resistance to bacterial pathogens in salmon and medaka (Arkoosh *et al*. 1998; Carlson *et al*. 2002). Together with the moderate toxicity levels measured in our study, these results point to adaptation driven primarily by indirect effects on physiology and immunity, rather than by strong, mortality-inducing selective pressures from pollutants.

### Adaptation to other environmental parameters

Aside from the associations with pollution, results from the Baypass core analysis revealed an abundance of global adaptive signals throughout the genome, with 162 regions showing evidence for positive selection. Interestingly, pollution-associated windows showed only limited overlap with these XtX windows (6.3%). This suggests that most loci under positive selection reflect responses to environmental pressures other than pollution. Two environmental factors, temperature and salinity, are plausible additional selective pressures in our study system that we did not account for. Given that *G. holbrooki* likely colonised Mediterranean regions from North-Eastern US populations near the Potomac River (Vera *et al*. 2016), climatic differences, particularly the higher temperatures in southern France, may represent a major driver of local adaptation. In addition, three populations in our dataset (MEJ, CAV and SAV) were sampled from brackish or saline habitats (with salinity ranging from 6 - 29.8 practical salinity units), and the species is also found in multiple saltwater sites across the region (E. Farcy, pers. comm.). It is therefore possible that an important driver of adaptation to the invaded Mediterranean habitats of *G. holbrooki* could be the response to temperature and/or salinity.

Thermal adaptation in fish is well described at the genetic level, with important genes from the heat shock protein family (*hsp*) differentially expressed in the short-term response to thermal stress (*e.g.,* Smith *et al*. 2013), or showing evidence of local adaptation in populations from distinct thermal environments (Chen *et al*. 2018). Similarly, adaptation to salinity in aquatic organisms relies on a set of well-known transmembrane proteins involved in the regulation of osmotic pressure. Genes coding for ion-transporters such as the V-type or Na/K ATPase are primary targets of selection during salinity transitions of aquatic organisms (Stern & Lee 2020, Velotta *et al*. 2022). Among the genes located within XtX peaks, we identified two loci on chromosomes 7 and 18 overlapping *hsp* genes (hsp12A and hsp13), but none associated with ion-transport genes. Other genes located at the intersections with the XtX windows covered a wide array of molecular functions, but no clear signal allowed us to wave in favour of adaptation to a particular environmental pressure. In that context, while the pollution-associated windows only represent 6.3% of the global genomic adaptive signal of our dataset, the specificity of the selection signal and the relevant biological processes involved suggest that persistent phenotypes have evolved in response to sublethal pollution cocktail exposure in the Mediterranean populations. Further characterization of the phenotypic variation in these populations through the implementation of reciprocal transplant experiments would be valuable to determine the traits under selection, both in response to pollution and other environmental parameters.

### Adaptation from reduced standing genetic variation?

Although we identified clear adaptive loci associated with pollution in our Mediterranean *G. holbrooki* populations, the age of these mutations and their frequency in ancestral populations is unknown. The detailed demographic history of *G. holbrooki* in Europe has yet to be described, but general patterns of invasion have been identified. Previous studies using mitochondrial and targeted SNP data (Vidal *et al*. 2010, Vera *et al*. 2016) have shown that southern European populations of *G. holbrooki* likely originated from the north-eastern USA (*i.e.,* near the Potomac River), and were further spread in the Mediterranean region by human activities. Organic and inorganic pollution is pervasive, and it is possible that source populations to our study populations have been exposed to similar cocktails of pollutants and that the adaptive loci we describe constitute standing genetic variation. Such a scenario aligns with the Anthropogenically Induced Adaptation to Invade (AIAI) hypothesis (Hufbauer *et al*. 2012), which posits that recent adaptation to human-disturbed habitats within the native range can pre-adapt populations and thereby enhance invasion success in similarly disturbed habitats abroad. Nonetheless, whereas some pollutants, such as metals linked to mining and coal industries, have long been present in aquatic ecosystems (Strzebońska et al. 2017), the use of PAHs, PCBs and PBDEs increased sharply only more recently, particularly from the second half of the 20th century onward (Heim & Schwarzbauer 2013). Similarly, the increase in diversity of molecules constituting pharmaceutical pollutants is a rather recent phenomenon (Wilkinson *et al*. 2022; Ortuzar *et al*. 2022), pointing towards the possibility that invasive European populations have been exposed to cocktails of pollution during their invasion, rather than prior to their introduction in the 1920s. Further investigation of *G. holbrooki*’s invasion history, based on broader sampling across both the native and invaded ranges, is needed to test the AIAI hypothesis, with particular emphasis on the age and frequency of the adaptive loci identified here within ancestral populations.

Although the detailed demographic history of our study populations remains relatively unknown (but see Vidal *et al*. 2011, Sanz *et al*. 2013), the European invasion of *G. holbrooki* almost certainly involved a drastic loss of genomic diversity (Vidal *et al*. 2011). Despite this severe bottleneck, we found that adaptive genomic regions show higher nucleotide diversity than the genomic background. This pattern points to rapid adaptation driven by standing genetic variation rather than de novo mutations, consistent with soft selective sweeps (Messer & Petrov 2013). Similar conclusions were drawn by Reid *et al*. (2016) study of adaptation to lethal levels of persistent organic pollutants in the Atlantic killifish (*Fundulus heteroclitus*). Specifically, their study showed that, while repeated selection on the same genes of the AHR pathway occurred across multiple populations of *F. heteroclitus*, multiple different genomic variants were also found to increase in frequency both in pollution-resistant and pollution-sensitive populations. Our results expand this body of work and show that adaptive evolution through soft selective sweeps fuelled by standing genetic variation can also occur in the face of sublethal exposure to pollutants, on a relatively short time scale (ca. 100 years).

## CONCLUSION

Our study provides important insights into the genomic bases of adaptation to multiple pollutants, and more generally into the genetics of adaptation during biological invasions. We show that populations persisting in polluted environments show specific changes in allele frequencies for genes involved in important biological processes, suggesting that selection for such allelic variants is at play in contaminated waters. We also show that adaptive genomic regions have higher nucleotide diversity than background genomic regions, reinforcing the idea that standing genetic variation in otherwise genetically depleted populations is important for rapid adaptation to novel environmental conditions during biological invasions. These results open important questions about how general this mechanism may be across taxa and stressors, and underline the need to integrate multifactorial pressures into future studies of adaptation in human-altered environments.

## Acknowledgements

The authors wish to thank the MGX-GENOMICS platform for sequencing; Bioinfo Genotoul https://doi.org/10.15454/1.5572369328961167E12 for computing facilities. We are grateful to A. Ducasse, for helping with the administration and funding management of the project; P. Nouhaud, T. Muller, M. Gautier and the GenPop group at CBGP for discussions regarding bioinformatic and statistical analyses.

## Authors’ contributions

AF analysed the data and led the writing of the manuscript. NM and EF sampled all populations and acquired pollution data. EBB, CR and NM performed DNA extraction. CP, AE and CR conceived the study. CP, AE, CR and EF secured funding for the study. All authors participated in editing and revising the manuscript.

## Data availability statement

All raw data used in this study will be deposited on the IFREMER data repository Dataref; Scripts necessary to replicate the analyses are publicly available on github (github/afraimout).

