## Supplementary material for "Genome-Environment Associations reveal shared and unique adaptive loci across multiple pollutants and populations of the eastern mosquitofish *Gambusia holbrooki*"

**Table of content**


| **Name** | **Page** |
|---|---|
| Figure S1. Manhattan plot of genome-wide FST | 2 |
| Figure S2. Nucleotide diversity between populations and genomic regions | 3 |
| Table S1. Samples information | 4 |
| Table S2. Results of the Gene Ontology enrichment analysis | 5 |

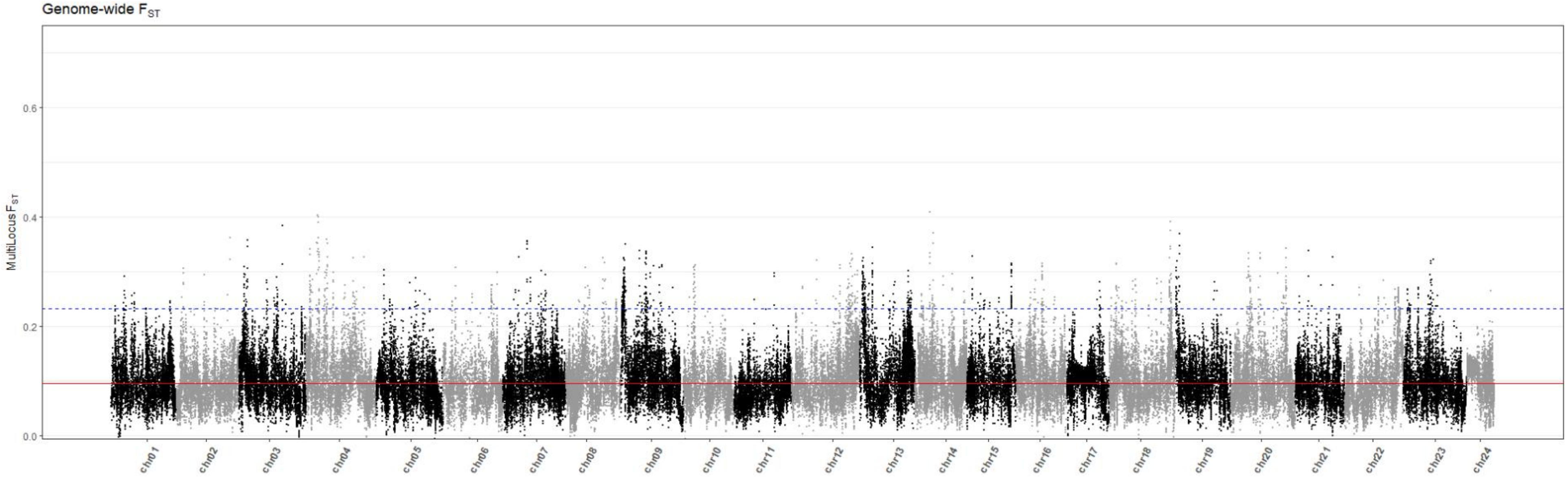


**Figure S1. Manhattan plot of genome-wide FST.** The multi-locus genome-wide FST calculated from *poolfstats* is shown along the 24 chromosomes (alternating black and gray dots). The median FST is shown by the red horizontal solid line and the top 1% of the distribution is shown by the blue horizontal dashed line.

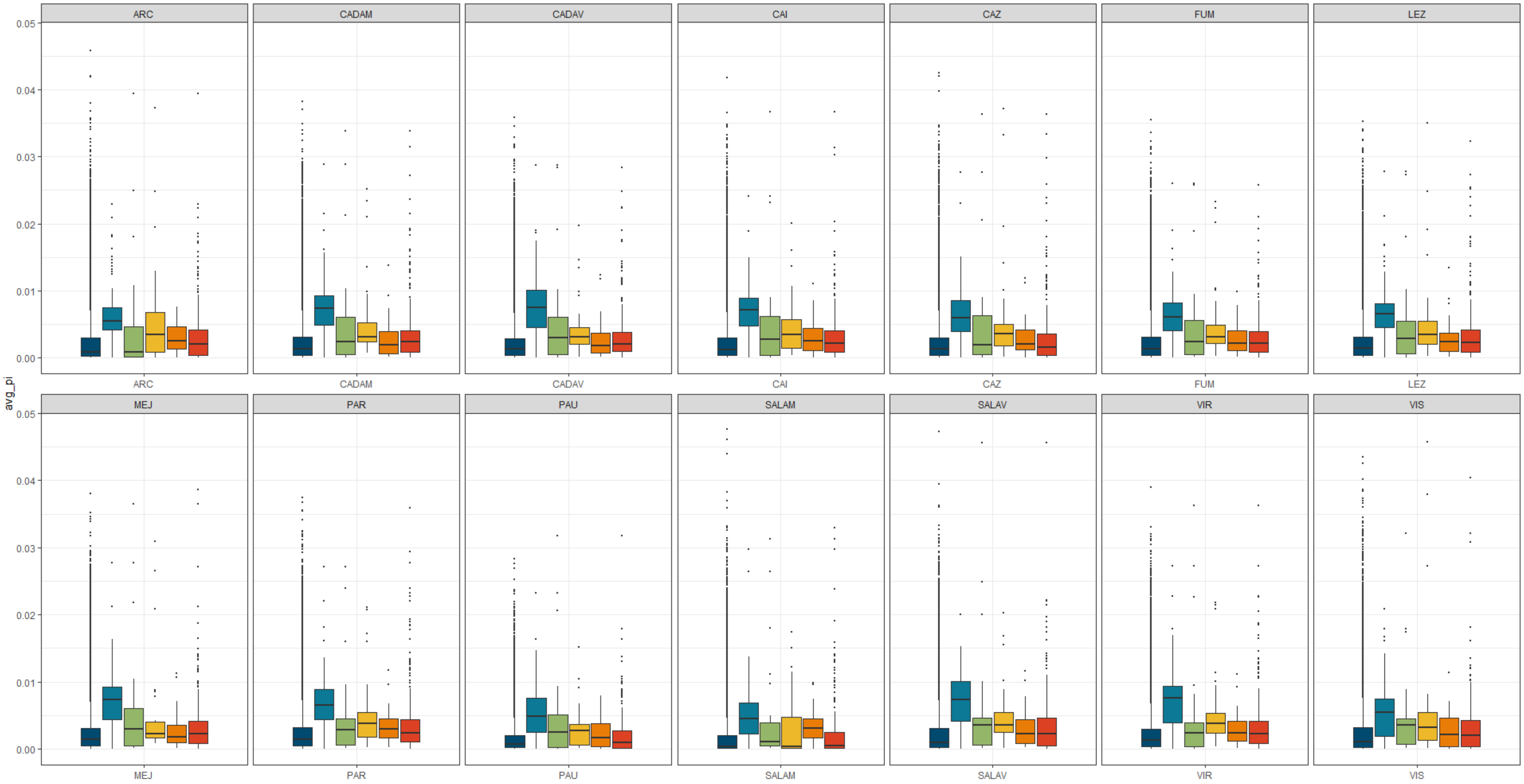


**Figure S2. Nucleotide diversity between populations and genomic regions.** The average nucleotide diversity (avg_pi) is shown for all categories of genomic windows: background (dark blue), lipophilic organics (blue), essential metals (green), non-essential metals (yellow), polar organics (orange) and XtX (red).

**Table S1. Samples information.** For each locality, the population code ("pop.code") is indicated along the GPS coordinates (XY; WGS84 format) and the number of individuals sequenced for the IndSeq dataset ("n.ind") or the PoolSeq dataset ("n.ind.pool"; and see *Methods* in the main text for the definition of both datasets.

| **pop.code** | **GPS coordinate X** | **GPS coordinate Y** | **n.ind** | **n.ind.pool** |
|---|---|---|---|---|
| ARC | 5.132 | 43.505 | 2 | 30 |
| CAM | 4.042 | 43.616 | 10 | 30 |
| CAV | 4.049 | 43.608 | 10 | 30 |
| CAI | 5.198 | 43.414 | 10 | 30 |
| CAZ | 3.708 | 43.763 | 10 | 30 |
| FUM | 4.682 | 43.513 | 10 | 30 |
| LEZ | 3.901 | 43.577 | 10 | 30 |
| MEJ | 3.9 | 43.542 | 10 | 30 |
| PAR | 3.898 | 43.572 | 10 | 30 |
| PAU | 3.869 | 43.632 | 10 | 30 |
| SAV | 4.018 | 43.595 | 2 | 30 |
| SAM | 4.002 | 43.609 | 2 | 30 |
| VIR | 4.075 | 43.637 | 10 | 30 |
| VIS | 4.214 | 43.607 | 2 | 30 |

**Table S2. Results of the Gene Ontology enrichment analysis.** GO.ID: the Gene Ontology identifier of the biological process; Term: the name of the biological process; Annotated: the number of genes annotated in the gene universe; Significant: the number of genes in the genomic window associated with pollution falling in that category; Expected: the number of genes expected to fall in this category following the null hypothesis of no enrichment. Rank in elimFisher: the rank of the GO term in the list based on the p-value from the "elim" Fisher test; classicFisher: the p-value for Fisher's exact test; elimFisher: the p-value fo the "elim" Fisher test; Pollutant: the type of pollutant associated with the genomic windows.

| **GO.ID** | **Term** | **Annotated** | **Significant** | **Expected** | **Rank in elimFisher** | **classicFisher** | **elimFisher** | **Pollutant** |
|---|---|---|---|---|---|---|---|---|
| GO:0045087 | innate immune response | 49 | 4 | 0.08 | 1 | 7.30E-07 | 3.10E-05 | lipophilic_organics |
| GO:0032461 | positive regulation of protein oligomerization | 1 | 1 | 0 | 2 | 0.0017 | 0.0017 | lipophilic_organics |
| GO:0032727 | positive regulation of interferon-alpha production | 1 | 1 | 0 | 3 | 0.0017 | 0.0017 | lipophilic_organics |
| GO:0032728 | positive regulation of interferon-beta production | 1 | 1 | 0 | 4 | 0.0017 | 0.0017 | lipophilic_organics |
| GO:0006260 | DNA replication | 53 | 2 | 0.09 | 5 | 0.0032 | 0.0032 | lipophilic_organics |
| GO:0060337 | type I interferon-mediated signaling pathway | 2 | 1 | 0 | 6 | 0.0033 | 0.0033 | lipophilic_organics |
| GO:0140374 | antiviral innate immune response | 2 | 1 | 0 | 7 | 0.0033 | 0.0033 | lipophilic_organics |

| | | | | | | | | |
|---|---|---|---|---|---|---|---|---|
| GO:0070534 | protein K63-linked ubiquitination | 5 | 1 | 0.01 | 8 | 0.0083 | 0.0083 | lipophilic_organics |
| GO:0010508 | positive regulation of autophagy | 6 | 1 | 0.01 | 9 | 0.0099 | 0.0099 | lipophilic_organics |
| GO:0070936 | protein K48-linked ubiquitination | 8 | 1 | 0.01 | 10 | 0.0132 | 0.0132 | lipophilic_organics |
| GO:0050728 | negative regulation of inflammatory response | 10 | 1 | 0.02 | 11 | 0.0165 | 0.0165 | lipophilic_organics |
| GO:0031348 | negative regulation of defense response | 12 | 1 | 0.02 | 12 | 0.0198 | 0.0198 | lipophilic_organics |
| GO:0050727 | regulation of inflammatory response | 16 | 1 | 0.03 | 13 | 0.0263 | 0.0263 | lipophilic_organics |
| GO:0032102 | negative regulation of response to external stimulus | 20 | 1 | 0.03 | 14 | 0.0328 | 0.0328 | lipophilic_organics |
| GO:0030514 | negative regulation of BMP signaling pathway | 16 | 2 | 0.03 | 1 | 0.00036 | 0.00036 | polar_organics |
| GO:0008203 | cholesterol metabolic process | 31 | 2 | 0.06 | 2 | 0.00139 | 0.00139 | polar_organics |

| | | | | | | | | |
|---|---|---|---|---|---|---|---|---|
| GO:0010874 | regulation of cholesterol efflux | 2 | 1 | 0 | 3 | 0.0037 | 0.0037 | polar_organics |
| GO:0044319 | wound healing, spreading of cells | 3 | 1 | 0.01 | 4 | 0.00554 | 0.00554 | polar_organics |
| GO:1902766 | skeletal muscle satellite cell migration | 3 | 1 | 0.01 | 5 | 0.00554 | 0.00554 | polar_organics |
| GO:0036342 | post-anal tail morphogenesis | 6 | 1 | 0.01 | 6 | 0.01105 | 0.01105 | polar_organics |
| GO:0040001 | establishment of mitotic spindle localization | 6 | 1 | 0.01 | 7 | 0.01105 | 0.01105 | polar_organics |
| GO:0048883 | neuromast primordium migration | 6 | 1 | 0.01 | 8 | 0.01105 | 0.01105 | polar_organics |
| GO:0048920 | posterior lateral line neuromast primordium migration | 6 | 1 | 0.01 | 9 | 0.01105 | 0.01105 | polar_organics |
| GO:0051293 | establishment of spindle localization | 7 | 1 | 0.01 | 10 | 0.01288 | 0.01288 | polar_organics |
| GO:0051653 | spindle localization | 7 | 1 | 0.01 | 11 | 0.01288 | 0.01288 | polar_organics |

| | | | | | | | | |
|---|---|---|---|---|---|---|---|---|
| GO:0048840 | otolith development | 12 | 1 | 0.02 | 12 | 0.02199 | 0.02199 | polar_organics |
| GO:0048881 | mechanosensory lateral line system development | 14 | 1 | 0.03 | 13 | 0.02561 | 0.02561 | polar_organics |
| GO:0048915 | posterior lateral line system development | 14 | 1 | 0.03 | 14 | 0.02561 | 0.02561 | polar_organics |
| GO:0048916 | posterior lateral line development | 14 | 1 | 0.03 | 15 | 0.02561 | 0.02561 | polar_organics |
| GO:0042074 | cell migration involved in gastrulation | 16 | 1 | 0.03 | 16 | 0.02922 | 0.02922 | polar_organics |
| GO:0048882 | lateral line development | 19 | 1 | 0.04 | 17 | 0.03461 | 0.03461 | polar_organics |
| GO:0048925 | lateral line system development | 19 | 1 | 0.04 | 18 | 0.03461 | 0.03461 | polar_organics |
| GO:1902850 | microtubule cytoskeleton organization involved in mitosis | 20 | 1 | 0.04 | 19 | 0.0364 | 0.0364 | polar_organics |
| GO:0035023 | regulation of Rho protein signal | 21 | 1 | 0.04 | 20 | 0.03819 | 0.03819 | polar_organics |

| | | | | | | | | |
|---|---|---|---|---|---|---|---|---|
| | transduction | | | | | | | |
| GO:0051028 | mRNA transport | 23 | 1 | 0.04 | 21 | 0.04176 | 0.04176 | polar_organics |
| GO:0002040 | sprouting angiogenesis | 26 | 1 | 0.05 | 22 | 0.04709 | 0.04709 | polar_organics |
| GO:0048598 | embryonic morphogenesis | 167 | 1 | 0.03 | 1 | 0.031 | 0.031 | metals_E |
| GO:0009790 | embryo development | 239 | 1 | 0.04 | 2 | 0.044 | 0.044 | metals_E |
| GO:0071277 | cellular response to calcium ion | 9 | 1 | 0 | 1 | 0.005 | 0.005 | metals_NE |
| GO:0006635 | fatty acid beta-oxidation | 20 | 1 | 0.01 | 2 | 0.0111 | 0.011 | metals_NE |
| GO:0019395 | fatty acid oxidation | 21 | 1 | 0.01 | 3 | 0.0116 | 0.012 | metals_NE |
| GO:0034440 | lipid oxidation | 22 | 1 | 0.01 | 4 | 0.0122 | 0.012 | metals_NE |
| GO:0009062 | fatty acid catabolic process | 28 | 1 | 0.02 | 5 | 0.0155 | 0.015 | metals_NE |
| GO:0072329 | monocarboxylic acid catabolic process | 33 | 1 | 0.02 | 6 | 0.0182 | 0.018 | metals_NE |
| GO:0030258 | lipid modification | 49 | 1 | 0.03 | 7 | 0.0269 | 0.027 | metals_NE |
| GO:0044242 | cellular lipid catabolic process | 59 | 1 | 0.03 | 8 | 0.0324 | 0.032 | metals_NE |

| | | | | | | | | |
|---|---|---|---|---|---|---|---|---|
| GO:0006631 | fatty acid metabolic process | 78 | 1 | 0.04 | 9 | 0.0427 | 0.043 | metals_NE |
| GO:0016042 | lipid catabolic process | 81 | 1 | 0.04 | 10 | 0.0443 | 0.044 | metals_NE |
| GO:0016054 | organic acid catabolic process | 89 | 1 | 0.05 | 11 | 0.0486 | 0.049 | metals_NE |
| GO:0046395 | carboxylic acid catabolic process | 89 | 1 | 0.05 | 12 | 0.0486 | 0.049 | metals_NE |
